## Supplemental Figures for "Cell cycle-gated feedback control mediates desensitization to interferon stimulation"

### Supplementary Figures

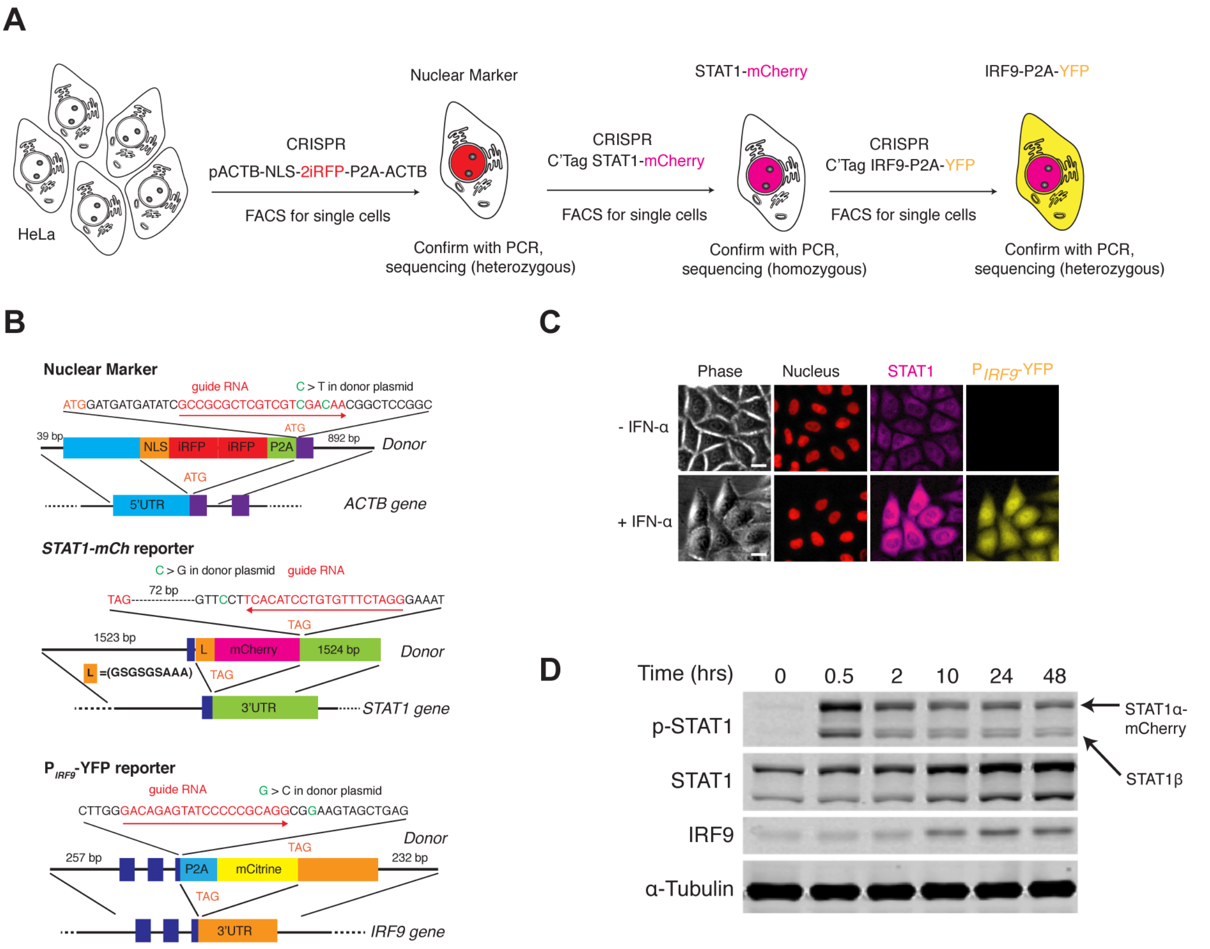


**Figure S1. Cell line construction and validation.** (A) Illustration of cell line construction steps. Full detail was described in Materials and Methods. Fluorescent reporters introduced and the targeted genes are shown. In each step, homogenous clones grew from single cells were carefully validated with PCR and sequencing to determine correct integration and homozygosity. Only one positive clone was selected to proceed with the next step. (B) Schematics of the *ACTB* (top), *STAT1* (middle) and *IRF9* (bottom) tagged loci. Sequences of the gRNA along with the recognition direction and the synonymous substitutions to avoid Cas9 recognition are shown. Targeted integration loci and the design of the donor DNA with indicated homology arms along with the inserts are also shown. (C) Representative images of the reporter cell line in response to IFN-α. Cells were treated without or with 100 ng/ml IFN-α for 48 hours. Scale bar: 20 μm. (D) Time course western blots showing the dynamics of phosphorylation (pY701) and expression of STAT1, and the dynamics of IRF9 expression in the reporter cell line. Cells were treated with 100 ng/ml IFN-α for indicated times, harvested and lysed for immunoblotting with indicated antibodies. The dynamics of the endogenous protein phosphorylation and expression, measured using immunoblotting, are similar to those measured using fluorescence microscopy (compare with Fig. 1C).

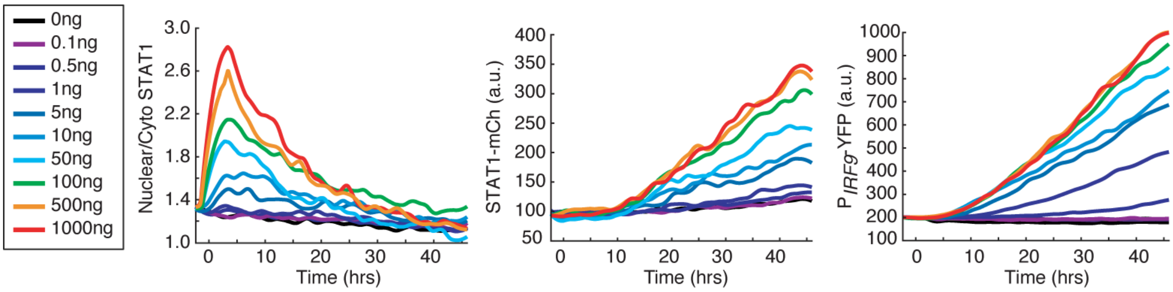


**Figure S2. Dose-dependent responses to IFN-α treatment.** Time traces of nuclear to cytoplasmic ratio for STAT1-mCherry, STAT1-mCherry fluorescence, and $P_{IRF9}$-YFP fluorescence in response to different concentrations (ng/ml) of IFN-α, as indicated. Averages of single cell traces were shown.

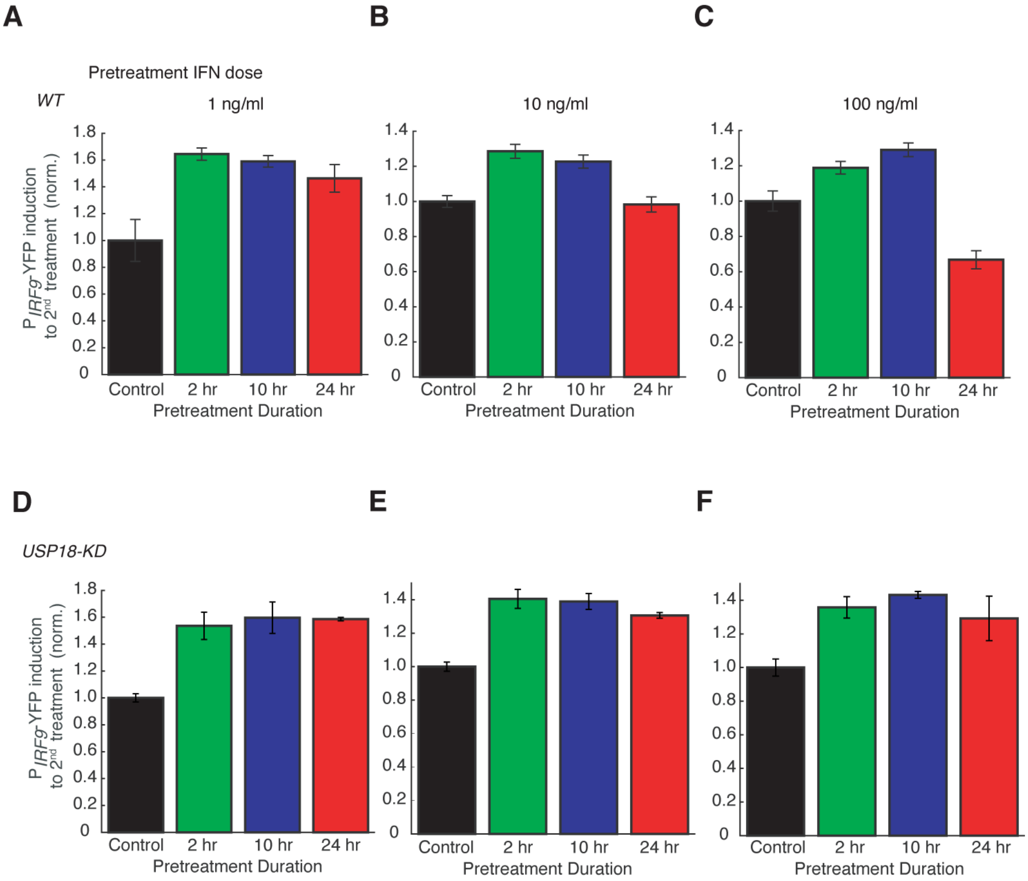


**Figure S3. Dose dependence of desensitization to IFN-α treatment.** Bar graphs showing the amounts of $P_{IRF9}$-YFP induction to the second IFN input (100 ng/ml) in WT (A - C) and USP18-KD (D – F), pretreated with different concentrations of IFN-α for different durations, as indicated. The results were normalized to the non-pretreatment condition (control).

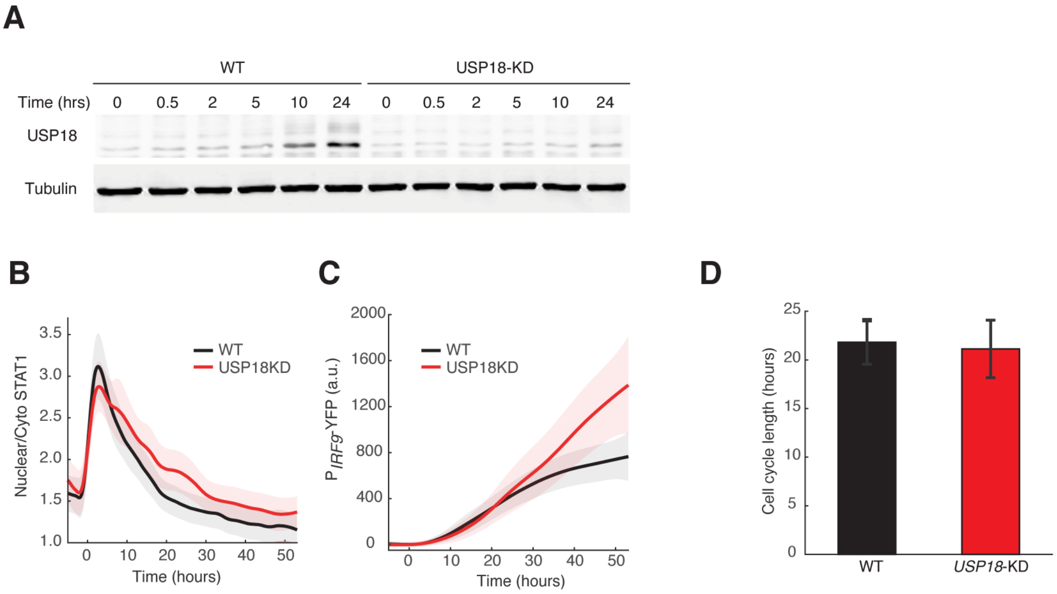


**Figure S4. Validation of the *USP18*-KD cell line.** (A) Western blots of USP18 expression in WT and *USP18*-KD cells. Cells were treated with 100 ng/ml IFN-α for indicated times, harvested and lysed for immunoblotting with the USP18 antibody. Time traces of nuclear to cytoplasmic ratio for STAT1-mCherry (B) and $P_{IRF9}$-YFP fluorescence (C) in WT (black) and *USP18*-KD (red) in response to IFN-α. Averages of single cell traces were shown. The shaded area represents ±SD. (D) Averaged cell cycle lengths of WT and *USP18*-KD. Cell divisions were identified in individual cells and the lengths between cell divisions were quantified. Error bars represent ±SD.

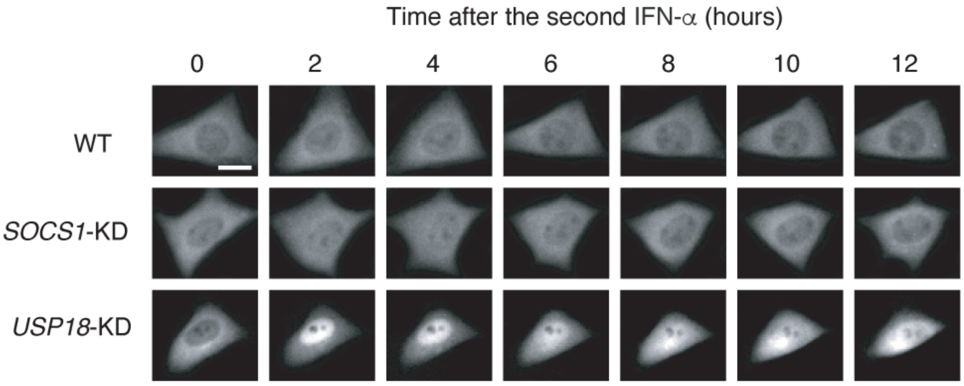


**Figure S5. SOCS1 does not mediate the desensitization of STAT1 nuclear translocation upon IFN stimulation.** Representative time-lapse images of STAT1 nuclear translocation upon the second IFN-$\alpha$ treatment in WT cells (top), *SOCS1*-KD (middle) and *USP18*-KD (bottom) cells. Cells were pretreated with 100 ng/ml IFN-$\alpha$ for 24 hours followed by 8 hours of break time and re-stimulated for 12 hours. Scale bar: 20 $\mu$m.

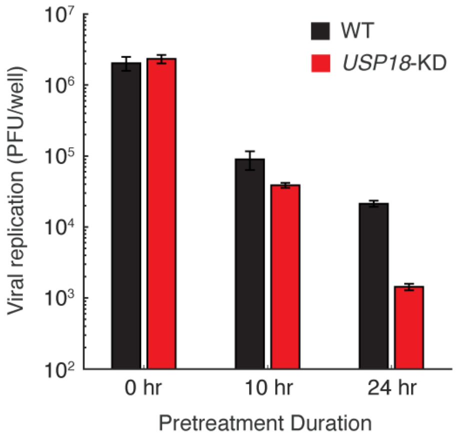


**Figure S6. Vesicular stomatitis virus (VSV) replication in WT and *USP18*-KD cells pretreated with different IFN-α durations.** Cell were pretreated with 100 ng/ml IFN-α for 0, 10 or 24 hours followed by 8 hours of break time and infected with 2500 PFU (plaque forming units) of VSV. After 18 hours, viral supernatant was collected and titered. Error bars represent ±SD, n=3. Note that the y-axis starts with 100 PFU/well as it is the limit of detection of our method.

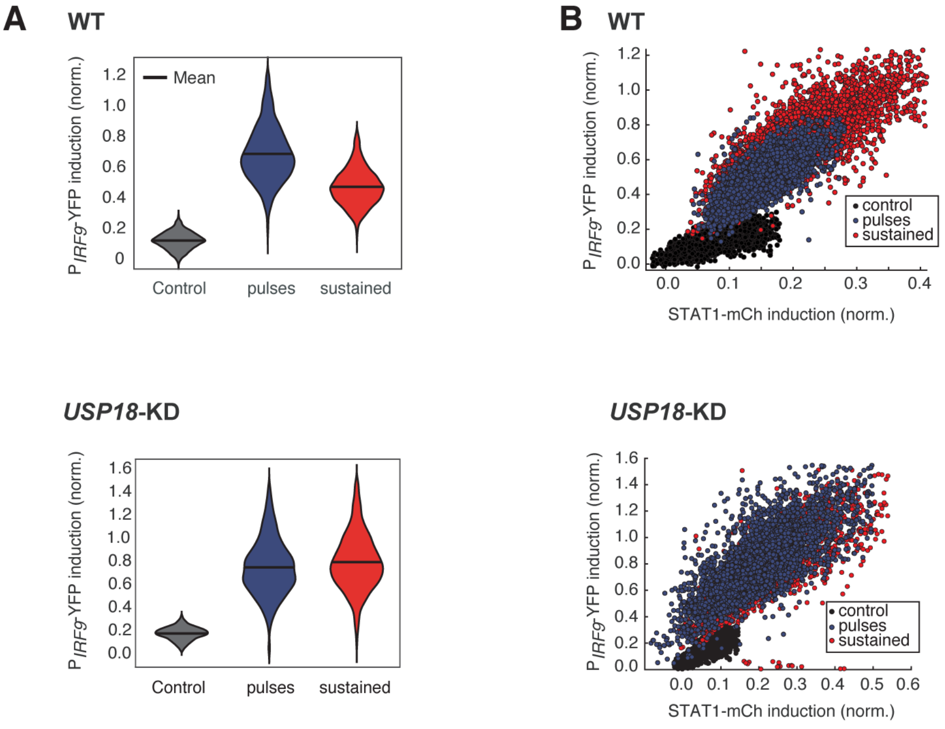


**Figure S7. Pulsatile IFN-α treatment induces higher ISG expression in single cells.** (A) The violin plots showing single-cell distributions of $P_{IRF9}$-YFP induction upon 5 x 8-hr pulsatile (blue) or 40-hr sustained (red) IFN-α treatments in WT (top) and *USP18*-KD (bottom) cells. (B) Scatterplots showing STAT1-mCherry versus $P_{IRF9}$-YFP induction in single cells in response to 5 x 8-hr pulsatile or 40-hr sustained IFN-α treatments in WT (top) and *USP18*-KD (bottom) cells. Fluorescent signals without IFN-α treatment were used as control.

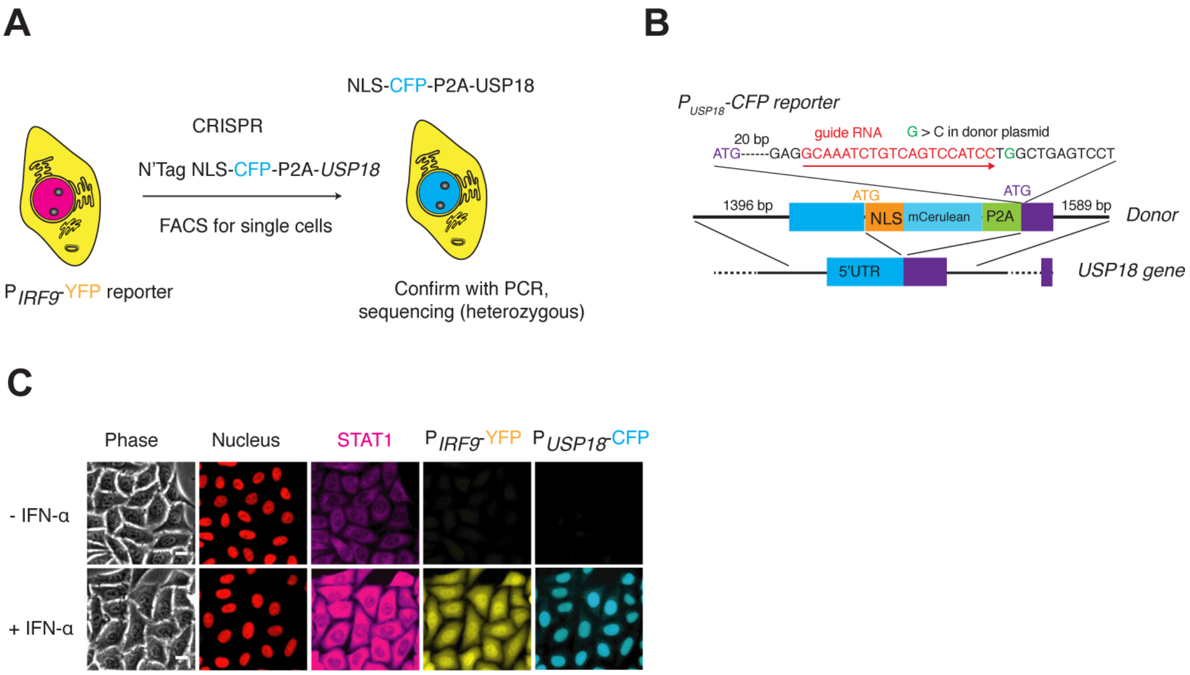


**Figure S8. Construction of the cell line with $P_{USP18}$-CFP reporter.** (A) Illustration showing the introduction of $P_{USP18}$-CFP into the dual reporter cell line in Fig. 1A. NLS-CFP-P2A coding sequence was inserted between the promoter and coding sequence of the *USP18* gene to generate a transcriptional reporter. (B) Schematic of *USP18* tagged locus. The sequence of the gRNA along with the recognition direction and the synonymous substitutions to avoid Cas9 recognition are shown. The design of the donor DNA with indicated homology arms along with the inserts are shown. (C) Representative images of the reporter cell line in response to IFN-$\alpha$. Cells were treated without or with 100 ng/ml IFN-$\alpha$ for 48 hours. Scale bar: 20 $\mu$m

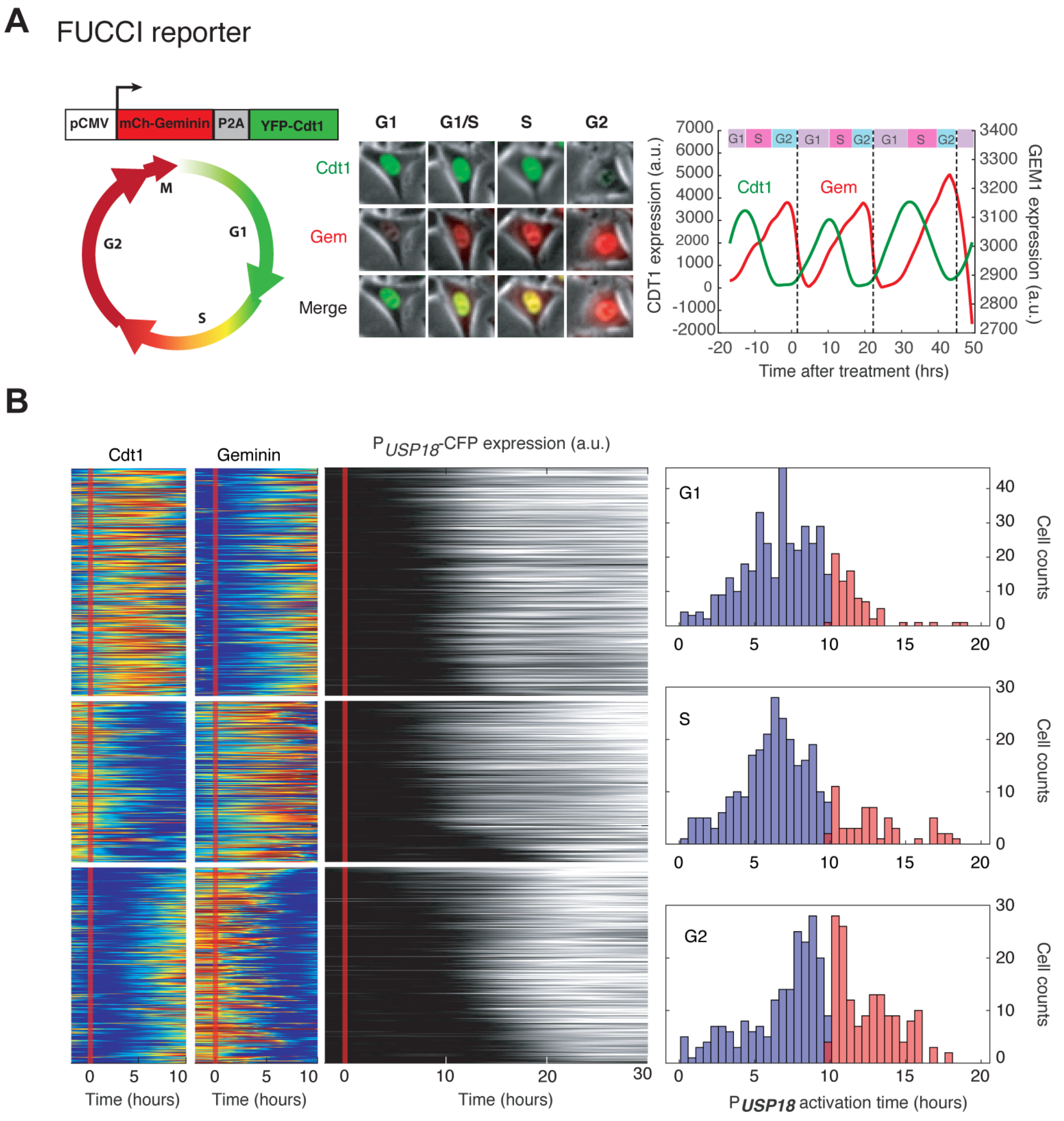


**Figure S9. Cell cycle-dependent *USP18* upregulation determined by the FUCCI reporter.** (A) Illustration of how the FUCCI reporter works. The fluorescent signals of chromatin licensing and DNA replication factor 1(Cdt1) and Geminin (Gem) proteins oscillate throughout a cell cycle to infer cell cycle phases. Middle: Images of a representative cell showing the Cdt1 and Gem level at different cell phases. Dynamics of Cdt1 and Gem signals in a single cell are shown along with the cell cycle phase inference. Dashed lines represent cell divisions. (B) Color maps of Cdt1, Gem and $P_{USP18}$-CFP expression in the same single cells. Each row represents the time trace of a single cell. Cells were grouped into G1 (n = 451), S (n = 388) and G2 (n = 325) based on Cdt1 and Gem signals (left) at the time of IFN-$\alpha$ addition. For each group, cells were sorted based on $P_{USP18}$-CFP activation time (middle). Right: Distributions of $P_{USP18}$-CFP activation times for each group.

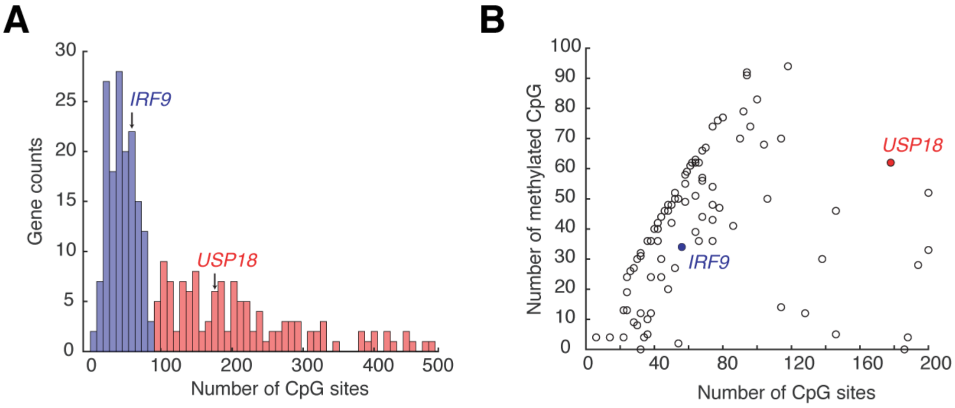


**Figure S10. ISG promoters contain a wide range of CpG site numbers and methylation levels.** (A) Histogram showing the numbers of CpG sites at ISG promoters. The promoter region is defined as 1000 bp upstream of the transcription start site. The data are collected from ENCODE database and the list of the 278 ISGs is from Interferome Database. (B) Scatterplot showing the numbers of methylated CpG versus of the numbers of CpG sites for ISG promoters. A CpG site is considered methylated if the methylation level is greater than 50% according to the bisulfide sequencing. Data are from ENCODE database for HeLa cells.

### Supplementary Movie

**Movie S1. Response of the dual reporter cell line to 100 ng/ml IFN-α treatment.** The image sequence was acquired for 50 hours in which the first two hours were before the addition of IFN-α. The NLS-2xiRFP nuclear marker is shown in red and is merged with phase images of the cells. STAT1-mCherry and $P_{IRF9}$-YFP in the same cells are also shown. Movie is shown at 50 frames per second.
